# Main sequence of human luminance-evoked pupil dynamics

**DOI:** 10.1101/2024.07.05.602284

**Authors:** Jonathan D Coutinho, Jeff Huang, Philippe Lefèvre, Gunnar Blohm, Douglas P Munoz

## Abstract

Pupil responses are commonly used to provide insight into visual perception, autonomic control, cognition, and various brain disorders. However, making inferences from pupil data can be complicated by nonlinearities in pupil dynamics and variability within and across individuals, which challenge the assumptions of linearity or group level homogeneity required for common analysis methods. In this study, we evaluated luminance evoked pupil dynamics in young healthy adults (N=10, 5:5 M:F, ages 19-25) by identifying nonlinearities, variability, and conserved relationships across individuals to improve the ability to make inferences from pupil data. We found a nonlinear relationship between final pupil diameter and luminance, linearized by considering the logarithm of luminance. Peak diameter change and peak velocity were nonlinear functions of log-luminance for constriction but not dilation responses. Across participants, curve fit parameters characterizing pupil responses as a function of luminance were highly variable, yet there was an across-participant linear correlation between overall pupil size and pupil gain (i.e. diameter change per unit log-luminance change). In terms of within-participant trial-by-trial variability, participants showed greater variability in final pupil size compared to constriction peak diameter change as a function of log-luminance. Despite the variability in stimulus-response metrics within and across participants, we found that all participants showed a highly stereotyped “main sequence” relationship between peak diameter change and peak velocity (independent of luminance). The main sequence relationship can be used to inform models of the neural control of pupil dynamics and as an empirical analysis tool to evaluate variability and abnormalities in pupil behaviour.

## Introduction

Pupil size is commonly measured to provide insight into visual perception, autonomic control, and cognition. Sensory information about visual luminance and defocus blur cause changes in pupil size and pupil size controls the amount of light entering the eye and the depth of field of the visual image (Barbur, 2004; Campbell and Green, 1965; Myers et al., 1990; Semmlow and Stark, 1973; Stark and Sherman, 1957). Clinical examination of pupil responses can be useful for differentiating and localizing neuropathology (Blumenfeld, 2010; Bremner, 2009; Kawasaki, 1999). Pupillography research has extensively catalogued correlations between pupil responses and cognitive phenomena including attention, arousal, memory, salience, and more (Ebitz and Moore, 2019; Einhäuser, 2017; Joshi and Gold, 2020; Mathôt, 2018; Strauch et al., 2022; Wang and Munoz, 2015). Given the multitude of sensory and- cognitive influences on pupil size, making inferences about neural activity from externally observable pupil behaviour can be challenging. This challenge is further complicated by nonlinearities and variability in pupil response dynamics.

There are two commonly described nonlinearities involved in luminance evoked pupil dynamics. First, both pupil size and firing rates of luminance neurons in the pretectal olivary nucleus (macaques) depend on the logarithm of luminance (Clarke et al., 2003; Ellis, 1981; Gamlin et al., 1995; Pong and Fuchs, 2000a, 2000b). Pupil size can be described by a linear or sigmoidal function of log-luminance, with the latter describing the saturation of pupil size at luminance extremes (Watson and Yellott, 2012). Second, pupillomotor responsiveness is nonlinearly modulated by instantaneous pupil size, where pupil size change per unit stimulus peaks at midrange pupil size. This effect can be observed from human behaviour (Semmlow et al., 1975), electrophysiological stimulation of the parasympathetic and sympathetic nerves in cats (Terdiman et al., 1971), and pharmacological experiments in isolated iris smooth muscle from rabbits (Yamaji et al., 2003). Modelling studies have related this size-dependent nonlinearity to the biomechanics of the iris smooth muscles (Fan and Yao, 2011; Semmlow and Stark, 1971). While these basic nonlinearities have been described, there remains limited quantification of potential nonlinear relationships in commonly used pupil metrics across a wide luminance range.

Another challenge for making inferences from pupil data is the limited characterization of variability in pupil responses within and across individuals. Within-participant variability includes changes in average stimulus-response relationships over time, and moment-by-moment changes in pupil size when environmental luminance and target distance are experimentally held constant (Stark et al., 1958). Within-participant variability also interacts with the size-dependent nonlinearity previously described, i.e. the magnitude of pupil variability depends on absolute pupil size (Stanten and Stark, 1966). Across-participant variability describes the differences between the average stimulus-response relationships between people. Previous studies have described the effects of age and light adaptation in explaining some variability of pupil responses (Watson and Yellott, 2012; Winn et al., 1994). Furthermore, one study focusing on across-participant variability in the pupil light reflex (with transient light flashes) found a wide range of pupil response variability between participants and correlated variability in pupil diameter change and velocity independent of age or mean pupil size (Bremner, 2012). Thus, there is a need for better quantification of within and across participant variability, particularly among young healthy adults (i.e. independent of aging and pathology).

The purpose of this study is to evaluate luminance evoked pupil dynamics in young healthy adults with a focus on quantifying response nonlinearities, variability, and conserved relationships across participants. We analyzed pupil responses from humans viewing step changes in greyscale luminance on a computer screen. We found a nonlinear relationship between final pupil diameter and stimulus luminance, linearized by considering the logarithm of luminance. However, other metrics such as peak change in diameter and peak velocity are still best described through nonlinear functions of log-luminance for constriction but not dilation responses. We quantify across-participant variability through the variability between linear regression parameters, finding an across-participant linear correlation between pupil gain and pupil size. Finally, despite the large degree of within-participant and across-participant variability in stimulus-response parameters (i.e. characterizing pupil response metrics as a function of the luminance stimulus), we describe a highly stereotyped main sequence relationship, i.e. a conserved nonlinear relationship between peak change in diameter and peak velocity for all subjects on a trial-by-trial basis. We introduce the use of pupil main sequence analysis as a research tool to better understand nonlinearities and variability in the neural control of pupil behaviour.

## Methods

### Participants

All experimental procedures were reviewed and approved by the Human Research Ethics Board of Queen’s University (protocol ID: PHYS-007-97). Ten participants (5:5 M:F) between the ages 19-25 years were recruited for this study. Our sample size was based on resource constraints rather than a priori power analysis, however we specifically set out to collect an equal sample size of male and female participants and to collect data from individual participants across two sessions on different days. All participants had normal or corrected-to-normal vision and provided written informed consent.

### Pupillography and Eye Tracking

Binocular pupil size and eye position were measured using an infrared, video-based eye tracker with a sampling rate of 500 Hz (Eyelink 1000, SR Research, Ottawa, Canada). Binocular pupil size measurements in video image pixels (proportional to pupil area) were scaled into millimeters diameter through a calibration procedure using false pupils of known size (Steiner and Barry, 2011; Wang et al., 2014).

Participants were seated comfortably in a darkroom with their heads restrained by a chinrest and headrest. Participants viewed a 17-inch LCD monitor with a resolution of 1280 x 1024 pixels and refresh rate of 60 Hz (Acer 1717). The monitor had a physical size of 33.9 cm x 27.1 cm and was placed at a 60.0 cm viewing distance, resulting in a viewing angle of 32° horizontal by 25° vertical. The configuration of the experimental setup (i.e. headrest, monitor, eye tracker camera) was held constant for all participants. Each experimental session began with a 9-point eye position calibration task (targets eccentricity ranging from ±14° horizontal and ±10° vertical from center) to ensure accuracy of eye position measurements.

### Behavioural Task

An experiment session had a duration of approximately 40 minutes total and consisted of 55 trials (a single trial is illustrated in Fig. 1). Each participant completed two sessions in total, with each session on separate days and differing only in the pseudorandomized sequence of luminance values presented in trials. Upon arriving at the lab for each session, participants confirmed their informed consent and were promptly seated in the darkroom to begin the calibration and experimental procedures within approximately 5-10 minutes (i.e., no enforced pre-experiment dark adaptation period, see Kelbsch et al., 2019). Participants were instructed to fixate the central dot and to minimize blinks during the experimental trials but that they may move their eyes and blink freely during the intertrial interval (7 s duration, 18.5 cd/m^2^ luminance). The central fixation stimulus had a diameter of 0.5° visual angle and a luminance of 18.5 cd/m^2^. We required central fixation to avoid potential measurement errors in pupil size due to eccentric eye position (Hayes and Petrov, 2016). Each trial consisted of a pseudorandomized sequence of 8 greyscale background luminance levels (ranging from 1 to 43 cd/m^2^ in steps of 7 cd/m^2^). Each background luminance value was presented for 5.2 seconds, resulting in trials of 41.6 s. This long stimulus presentation duration was selected to accurately evaluate pupil dynamics considering the slow time course of dilation responses. Pupil responses to the onset of the first luminance value in a trial were not analyzed to avoid confounding from the intertrial interval.

**Figure 1.**
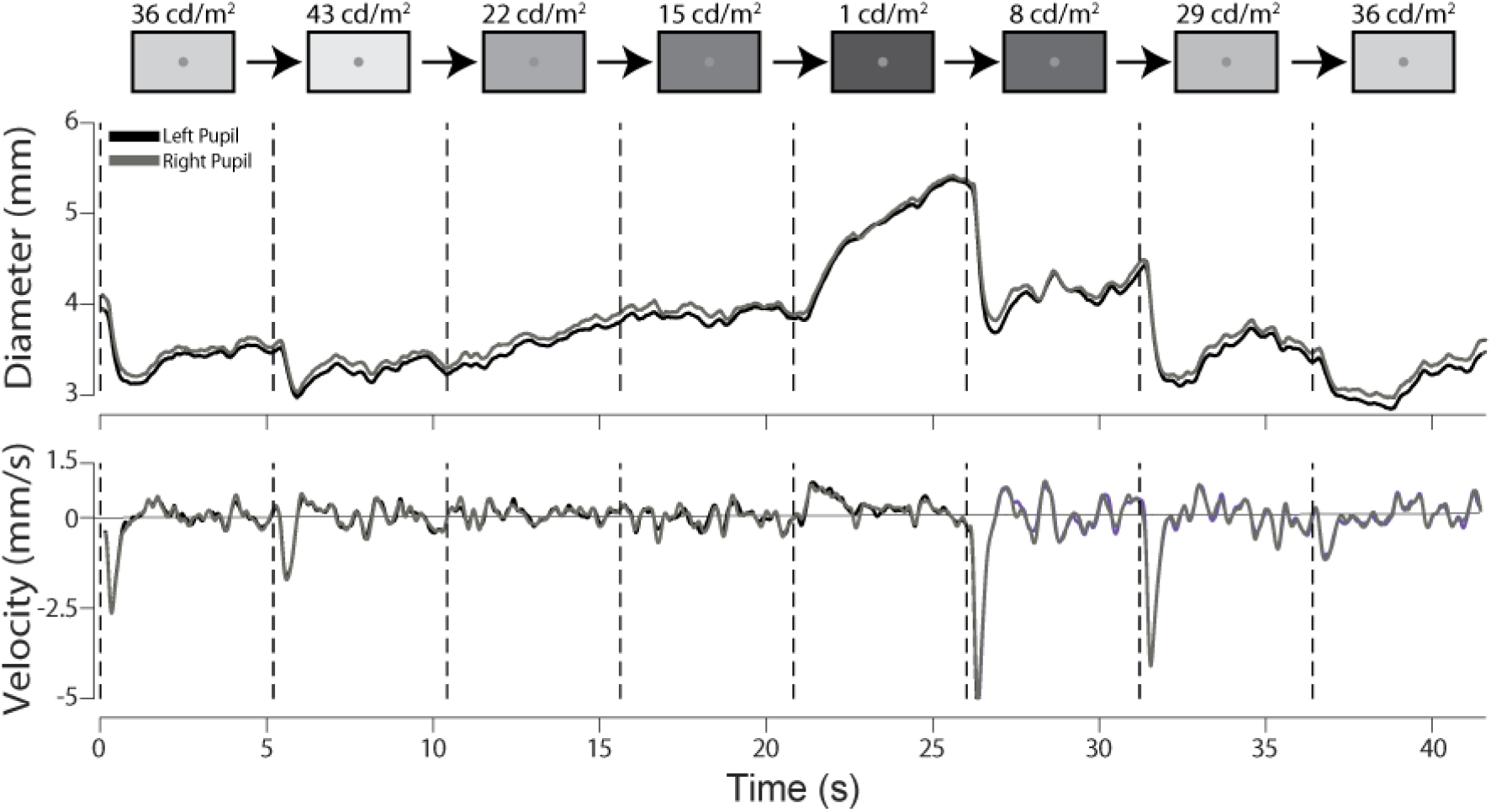
Example of a single trial. Participants centrally fixated a small dot (0.5° diameter, 18.5 cd/m^2^ luminance) while the background luminance stepped through a pseudorandom sequence of eight values (screen subtending 32° x 25° viewing angle, luminance values ranging from 1 to 43 cd/m^2^). The timing of luminance transitions is illustrated in black dashed vertical lines, with each luminance value presented for 5.2 seconds. Bilateral pupil diameter (mm) and pupil velocity (mm/s) are plotted over time for a full trial (41.6 seconds).

## Data Analysis

Our data consisted of time-series values of screen luminance (calibrated into cd/m^2^), pupil size (calibrated into millimeters diameter), and eye position (calibrated into 2D position in ° visual angle). These time-series data are recorded as trial segments corresponding to a pseudorandom sequence of 8 luminance values held constant for 5.2 seconds each (41.6 s total per trial). Data was segmented into single transitions, with a total of (2 pupils) x (7 transitions) x (55 trials) x (2 sessions) = 1540 transition segments per participant. Data were preprocessed to check for data loss (potentially corresponding to blinks) or deviation in eye position greater than 2°, resulting in the exclusion of 37% of the total 15,400 single transition time-series segments across participants. This conservative data inclusion strategy avoided the need for model-based pupil size corrections (e.g. Hayes and Petrov, 2016; Kret and Sjak-Shie, 2019; Mathôt et al., 2018). To better estimate instantaneous changes in pupil diameter (i.e. pupil velocity), data was filtered using autoregressive filtering with a cutoff frequency of 10 Hz. Fig. 2illustrates the pupil metrics extracted for each luminance transition: baseline pupil diameter (calculated from the 250 ms interval before luminance transition), final diameter (calculated from 4.90 to 5.15 s after luminance onset), peak diameter change, total diameter change, and peak velocity. Additionally, we computed the average time series pupil response to each (initial, final) luminance condition to compare across-participant variability, averaging together both sessions and both left and right pupils (see Supplementary Table 1 for comparison of bilateral pupil size across sessions). Mean pupil metrics were then calculated from the mean time series. All data analyses were performed using MATLAB 2019b (MathWorks, Inc., Natick, Massachusetts, USA). The data is available on https://osf.io/vrch6/ and all analysis code is available on https://github.com/blohmlab/PupilMainSequence.

**Figure 2.**
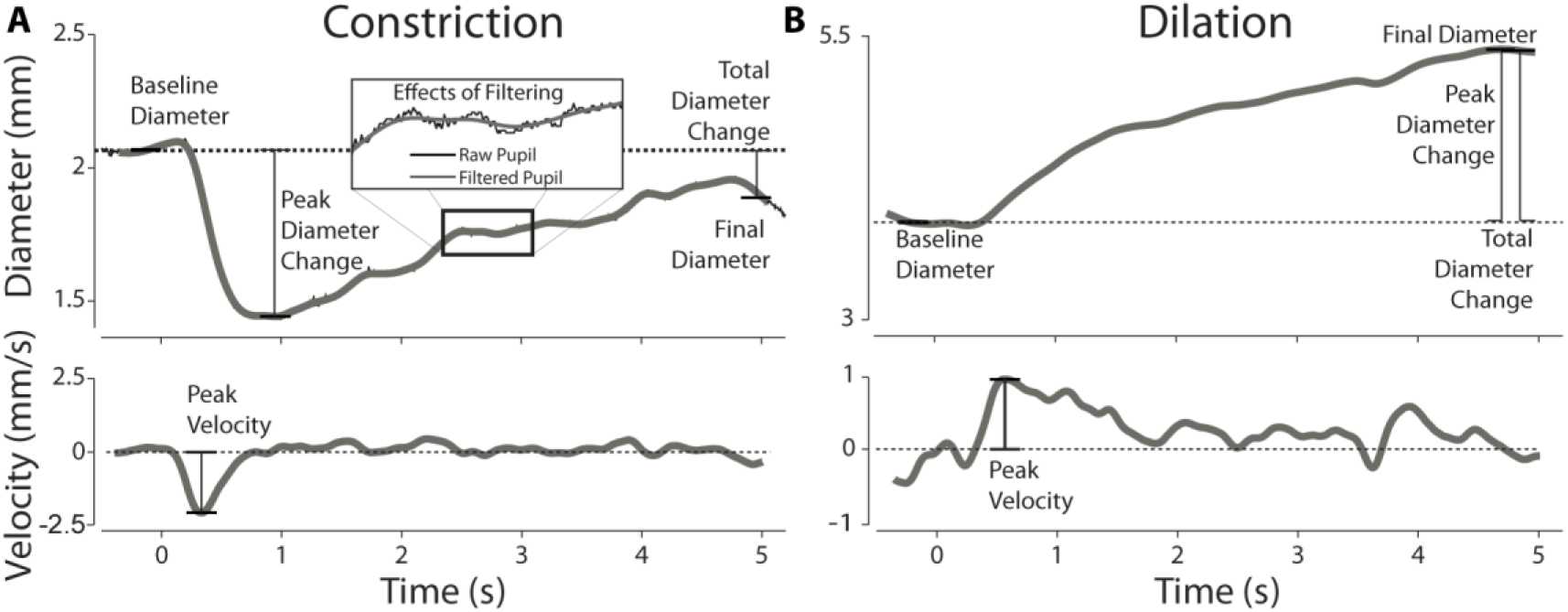
Examples of pupil response dynamics and metrics of interest during single luminance transitions. The luminance transitions occur at time=0. The inset in A illustrates the effects of autoregressive filtering (10 Hz cutoff frequency) on diameter data to subsequently calculate velocity from the filtered traces. The dashed horizontal line traces baseline pupil diameter and zero velocity on their respective axes. Note the difference in y-axis scales between the constriction and dilation responses. (A) Example of a constriction response, illustrating unilateral pupil diameter and velocity over time and associated metrics of interest. (B) Example of a dilation response, illustrating pupil diameter and velocity over time and associated metrics of interest.

## Results

The purpose of this experiment was to quantify luminance evoked pupil responses in humans. An example of a single trial is illustrated in Fig. 1, where bilateral pupil size is measured during step changes in large-field greyscale luminance. We analyzed the metrics characterizing steady state pupil behaviour (i.e. baseline pupil diameter, final pupil diameter, and total diameter change), and metrics characterizing the transient dynamics of pupil responses (i.e. peak diameter change and peak velocity). These behavioural metrics are illustrated in Fig. 2.

For each participant, final pupil diameter decreased with increasing luminance (Fig. 3). The relationship between luminance and pupil diameter was nonlinear and was characterized by an exponential curve-fit (Fig. 3A). When considering the luminance stimulus magnitudes on a logarithmic scale, the relationship was linear (Fig. 3B). This logarithmic relationship suggests that the pupil responds to the ratio of luminance change rather than the absolute change in luminance, further demonstrated by the linear relationship between changes in pupil diameter and changes in log-luminance (Fig. 3C). Thus steady-state pupil size for each individual participant was well characterized by a linear relationship between pupil diameter and log-luminance, yet there was a large degree of within-participant trial-by-trial variability in final pupil diameter and across-participant variability in the parameters of this linear relationship.

**Figure 3.**
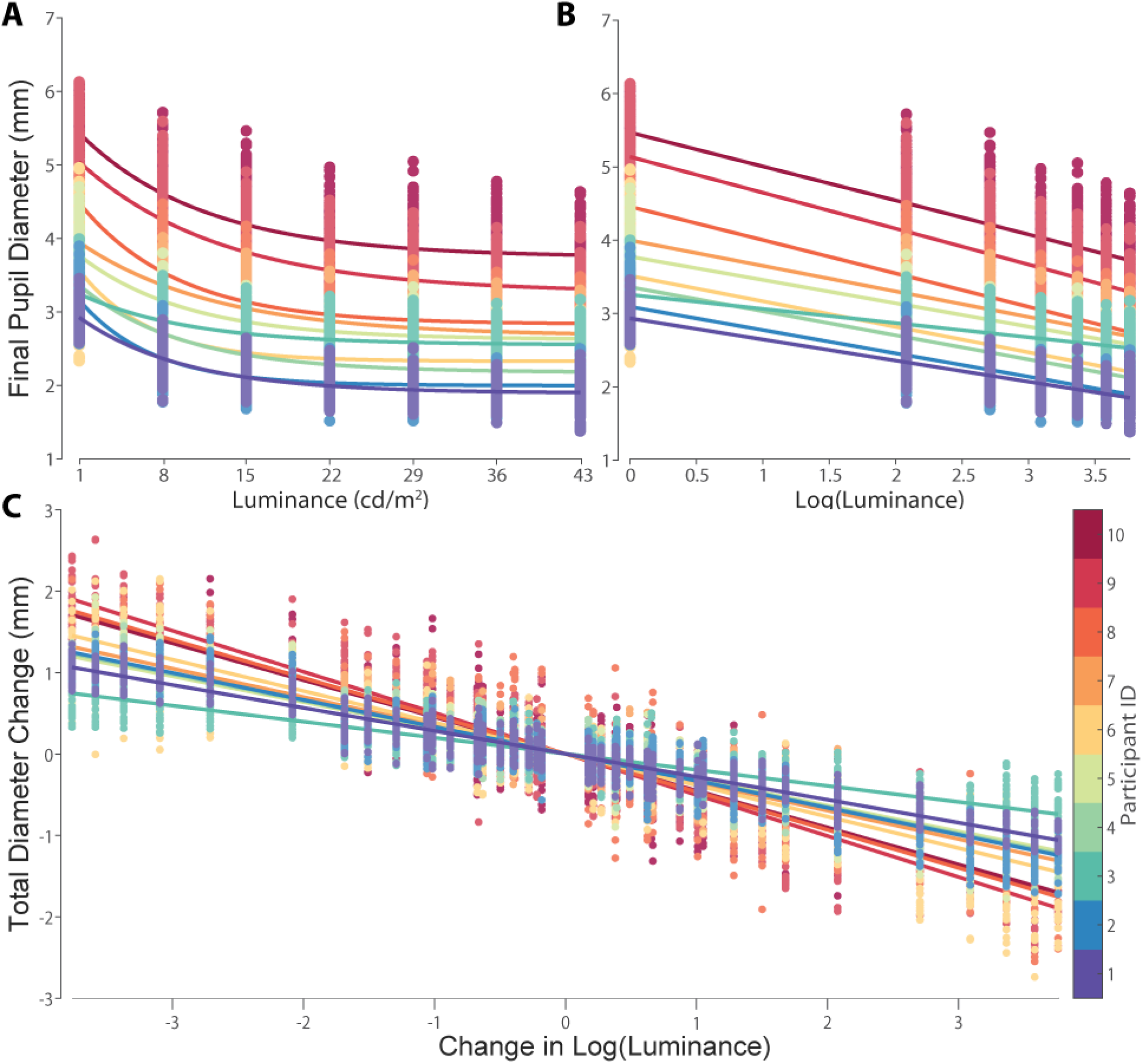
Transition-by-transition steady-state pupil diameter vs luminance. Data from individual participants are plotted in different colors, with colors sorted according to participants’ average baseline pupil diameter in the darkest condition (1 cd/m^2^). A) Final pupil diameter (time averaged from the last 250 ms of each luminance stimulus) was nonlinearly correlated with stimulus luminance for each subject, fit with an exponential function (y=a_1_*exp(-x/a_2_) + a_3_). B) Final pupil diameter vs log-luminance, fit by a linear function (y=a_1_*x+a_2_). C) Total diameter change vs change in log-luminance, fit by linear proportionality (y=a_1_*x).

The variability across participants in the relationship between pupil diameter and log-luminance can be numerically characterized through comparison of each subject’s linear curve-fit parameters.Specifically, this characterizes two parameters corresponding to the slope (pupil gain) and offset (fit diameter at log-luminance=0) for individual subjects. Figure 4 illustrates the linear correlation between these parameters across participants, demonstrating that participants with larger pupils also show larger pupil gain (i.e., larger changes in pupil diameter per log-unit change in luminance). This finding should be contrasted with previous descriptions of within-subject pupil-size dependent gain nonlinearity (e.g. Semmlow et al., 1975), which has been related to instantaneous pupil size and muscle-level nonlinearities. Instead, here we describe across-participant variability in average pupil gain as a function of average pupil diameter.

**Figure 4.**
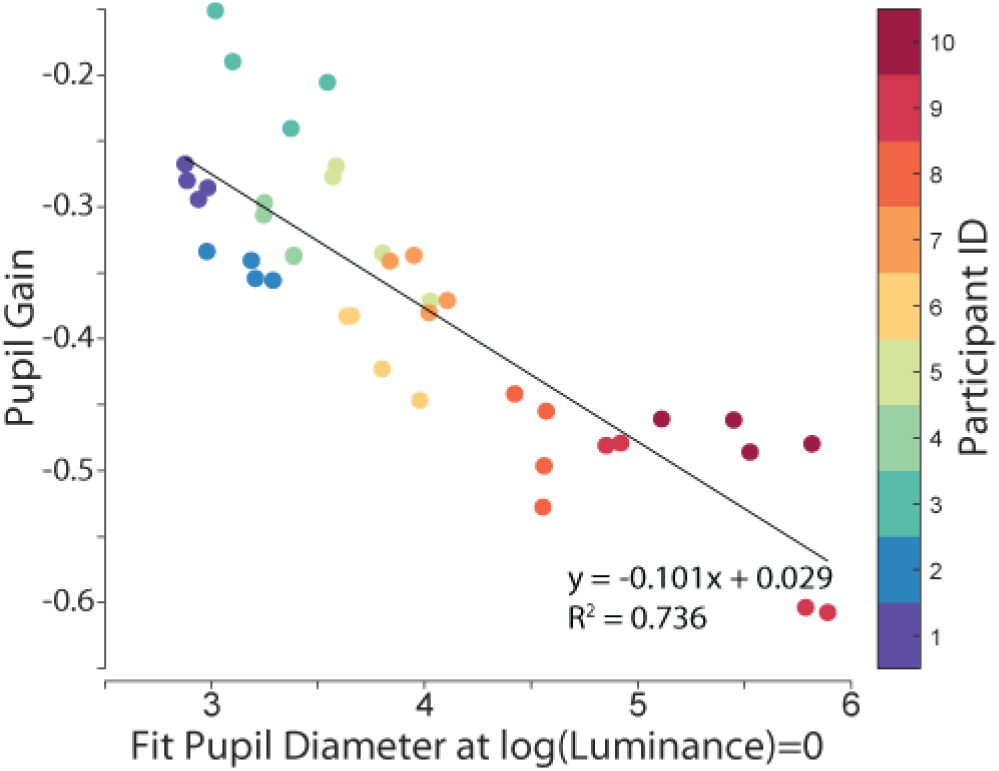
Across participant variability in linear regression parameters characterizing the relationship between final pupil diameter and log-luminance. Pupil gain, plotted on the y-axis, describes the amount of total pupil diameter change per log-unit change in luminance (i.e. the slope of the linear regression between final pupil diameter and log-luminance, Fig. 3B). The x-axis plots the fit value of pupil diameter when log(luminance)=0 or when luminance equals 1 cd/m^2^ (i.e. the y-intercept of the linear regression of final pupil diameter vs log-luminance, Fig. 3B). Thus, the values on the x-axis reflect a prediction for pupil diameter based on all luminance values, while the order of Participant ID color is based on the mean final pupil diameter at log-luminance=0 independent of pupil size at other luminance values. Each data point corresponds to a single eye during a single session (i.e. four data points per participant from bilateral pupillography across two sessions). The black solid line represents the linear fit across all participants.

Considering the transient dynamics of pupil responses, there were clear qualitative differences between the time-course of constriction vs dilation responses. Fig. 5 illustrates constriction and dilation dynamics to large and small magnitude log-luminance changes. Fig. 5A illustrates pupil responses for a single subject with an initial luminance value of 8 cd/m^2^ switching to either 43 cd/m^2^ (black lines) or 1 cd/m^2^ (orange lines, matching their Participant ID color). These examples illustrate constriction and dilation responses (respectively) to large magnitude changes in log-luminance. Note that constriction responses display larger magnitude peak velocity and earlier peak velocity times. Consequently, constriction responses also show large peak diameter changes early after the transition, overshooting their final values, while dilation responses continue to slowly dilate throughout. Fig. 5B plots the mean pupil response for the same conditions across all participants, showing similar trends for each participant.

**Figure 5.**
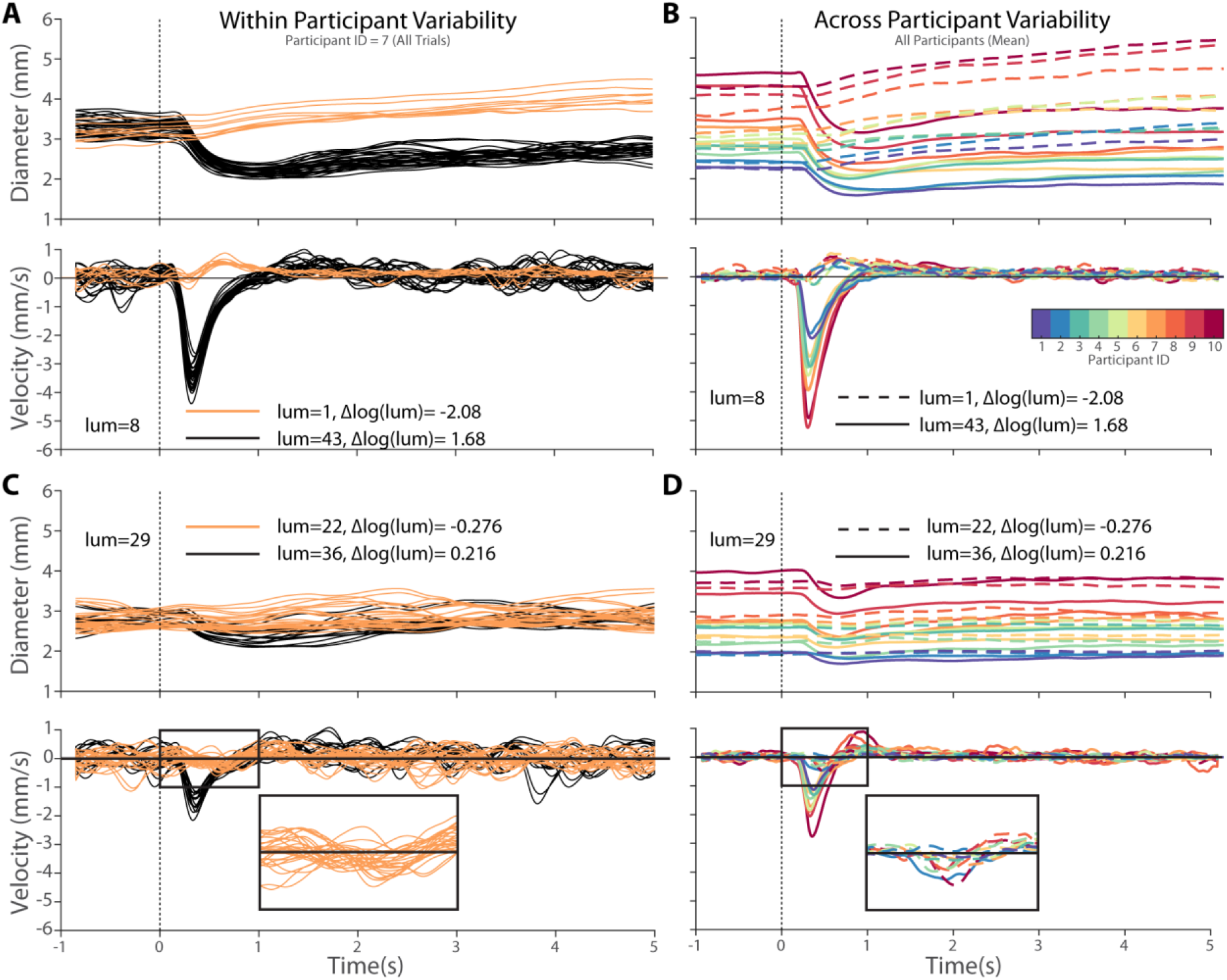
Within-participant variability in single transition pupil dynamics. **(A**,**C) and across-participant variability in mean pupil dynamics (B**,**D)**. Each row depicts luminance transitions beginning with the same value then increasing or decreasing in similar magnitude log-luminance steps. A) Single trial examples illustrating pupil diameter and velocity over time during large-magnitude luminance increase (from 8 cd/m^2^ to 43 cd/m^2^, 1.68 log-units and from 8 cd/m^2^ to 1 cd/m^2^, -2.08 log-units). B) Mean pupil diameter and velocity over time for the same luminance transition conditions for each participant. C) Single trial examples, illustrating pupil diameter and velocity over time during a small change in log-luminance (from 29 cd/m^2^ to 22 cd/m^2^, -0.276 log-units and from 29 cd/m^2^ to 36 cd/m^2^, 0.216 log-units).D) Mean pupil diameter and velocity over time for the same small-magnitude luminance changes for each participant. The inset in C,D illustrates pupil velocity from 0 to 1 s in the decreasing luminance condition at 2x magnification in both x-scale and y-scale, highlighting the negative velocity associated with the paradoxical constriction response.

Fig.5C illustrates pupil responses for a single participant during small magnitude changes in log- luminance, showing different qualitative features. Notably, during small decreases in luminance we observe occurrences of initial pupil constriction (Fig. 6C inset). This paradoxical constriction response has a different magnitude and timing compared to constriction responses to small increases in luminance: it was more similar in timing to the positive dilation velocity observed in large magnitude luminance decreases (Fig. 5A). Fig. 5D illustrates the averaged pupil response for each participant to the same small magnitude log-luminance transitions, again showing qualitatively similar across-participant responses.Thus, the transient dynamics of pupil responses show clear asymmetries between constriction and dilation and seem to nonlinearly depend on the magnitude of log-luminance change.

**Figure 6.**
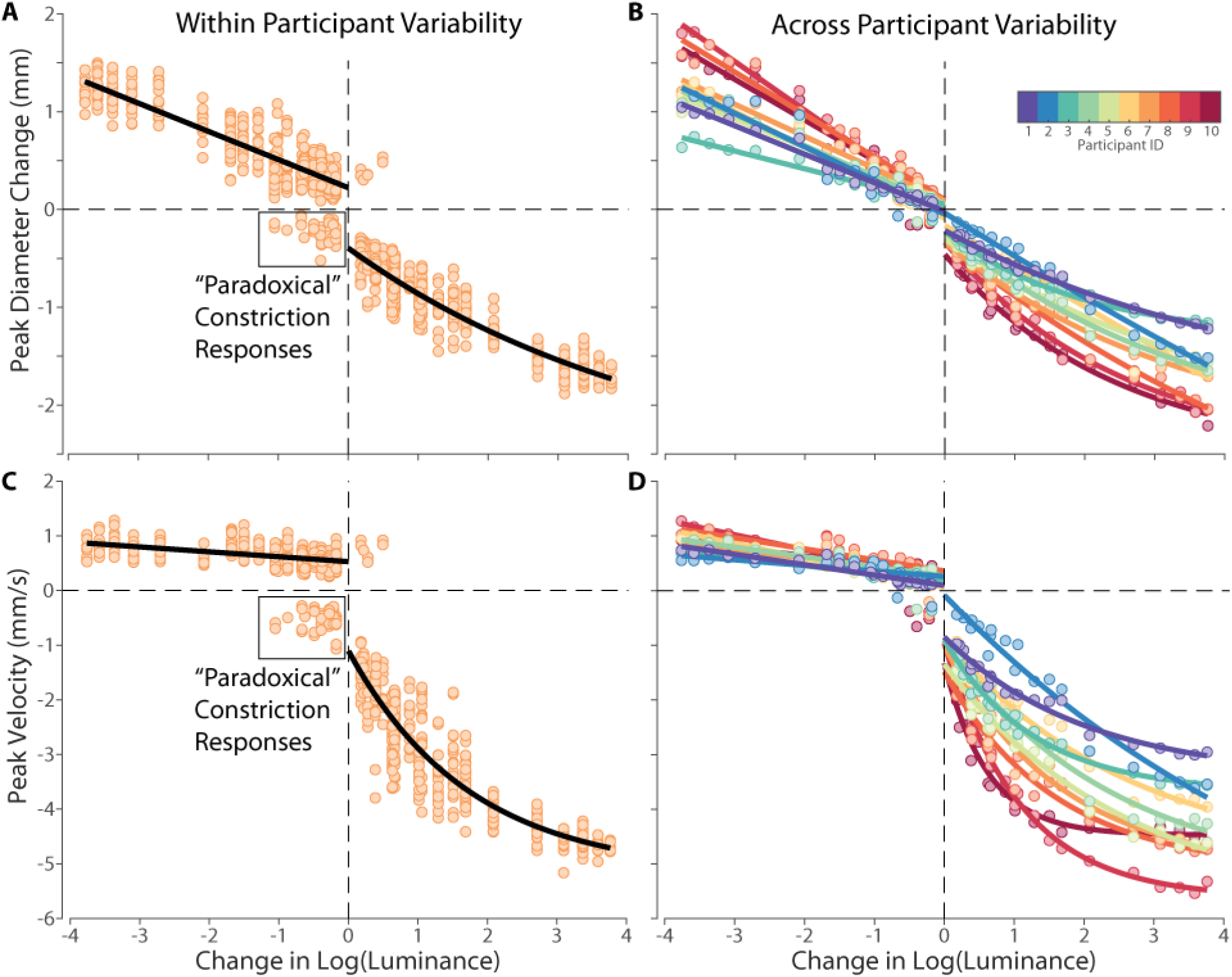
Transient pupil metrics: peak diameter change and peak velocity vs change in log-luminance. For each participant, dilation responses were fitted with a linear function (y = a_1_*x + a_2_) and constriction responses with an exponential function (y=a_1_*exp(x/a_2_) + a_3_). A) Transition-by-transition peak diameter change vs change in log-luminance from a single participant, highlighting within-participant variability. B) Peak diameter vs change in log-luminance for average pupil dynamics for each (initial, final) luminance condition for all participants, highlighting across-participant variability. C) Transition-by-transition peak velocity vs change in log-luminance for a single participant. D) Peak velocity vs change in log-luminance for average pupil dynamics for each (initial, final) luminance condition across participants.

We quantified the transient dynamics of pupil responses through the peak diameter change and peak velocity metrics. Figure 6 illustrates peak diameter change as a function of change in log-luminance. For decreases in log-luminance, pupil peak diameter changes generally followed a linear trend except for small magnitude log-luminance changes, where instead the initial paradoxical constriction often produced larger diameter changes than subsequent dilation. For increases in log-luminance, we observed a nonlinear relationship, reasonably fit with an exponential decay function characterizing a large initial value followed by saturation. The same trends were observed when considering peak velocity, illustrated in Fig. 6C,D. For small decreases in log-luminance, we observed many occurrences of greater initial constriction velocity than subsequent dilation peak velocity. Generally, dilation peak velocity was much lower than constriction peak velocity. Thus, even after considering luminance on a logarithmic scale, constriction peak diameter change and peak velocity involve further nonlinearities.

The timing of transient response metrics is illustrated in Fig. 7, grouping trials with increasing vs decreasing luminance. We observed similar trends in transient response timing across all participants. When luminance increases, the timing of peak diameter changes (Fig. 7A) typically occurs between 0.5 to 1.5 seconds after an increasing step in luminance. On the other hand, the timing of peak diameter change after decreasing luminance steps was highly variable, with a large proportion of responses showing continued dilation 5 seconds after the luminance transition. The timing of peak velocity (Fig. 7B) was highly regular for trials with increasing luminance, between 0.25 to 0.5 seconds. Trials with decreasing luminance showed much more variability in peak velocity timing between 0.5 to 1.0 seconds. Recall that dilation velocities are much lower than constriction velocities and the common occurrence of oscillations in pupil responses (Fig. 5A,C), leading to a spread of identified dilation peak velocity times over the time course of the trial. Thus, constriction transient response metrics are much more time-locked to the luminance transition and identifiable than dilation.

**Figure 7.**
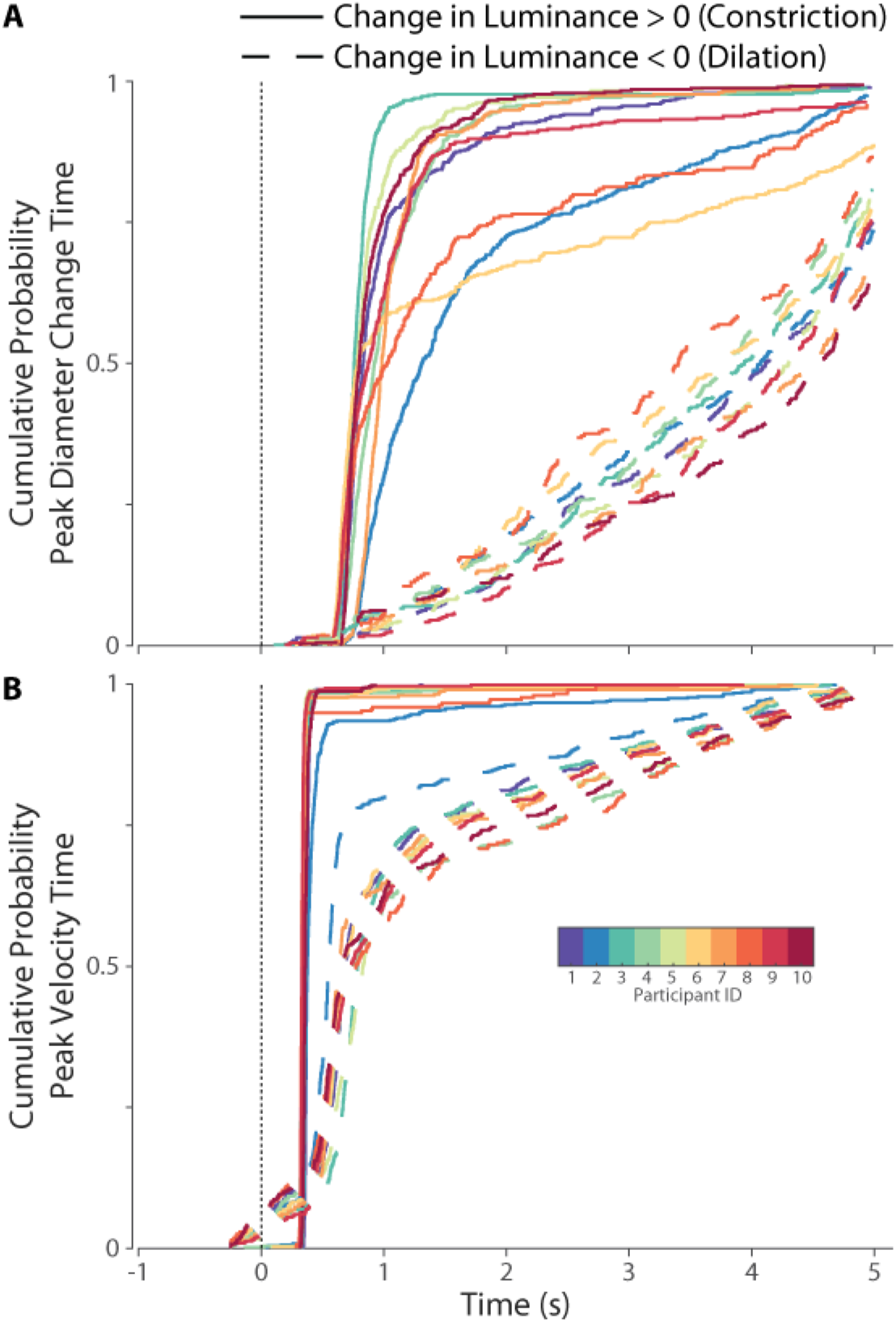
Cumulative probability distributions describing the timing of peak diameter change and peak velocity across participants. **A)** Cumulative probability describing the timing of peak diameter change for trials with increasing luminance (solid lines) or decreasing luminance (dashed lines). **B)** Cumulative probability distribution describing the timing of peak velocity for trials with increasing luminance (solid) or decreasing luminance (dashed).

To further evaluate within-participant trial-by-trial variability, we compared the variability in final pupil diameter to the variability in constriction peak diameter change (Fig. 8). We specifically consider constriction peak diameter change because of the timing regularity relative to the luminance transition, while final pupil diameter may include additional sources of variability accumulating over time. For all participants, we found greater trial-by-trial variability in final diameter compared to constriction peak diameter change. Thus, we observe more within-participant variability in final pupil size compared to transient constriction dynamics, suggestive of accumulated variability from uncontrolled parameters (i.e. unrelated to luminance).

**Figure 8.**
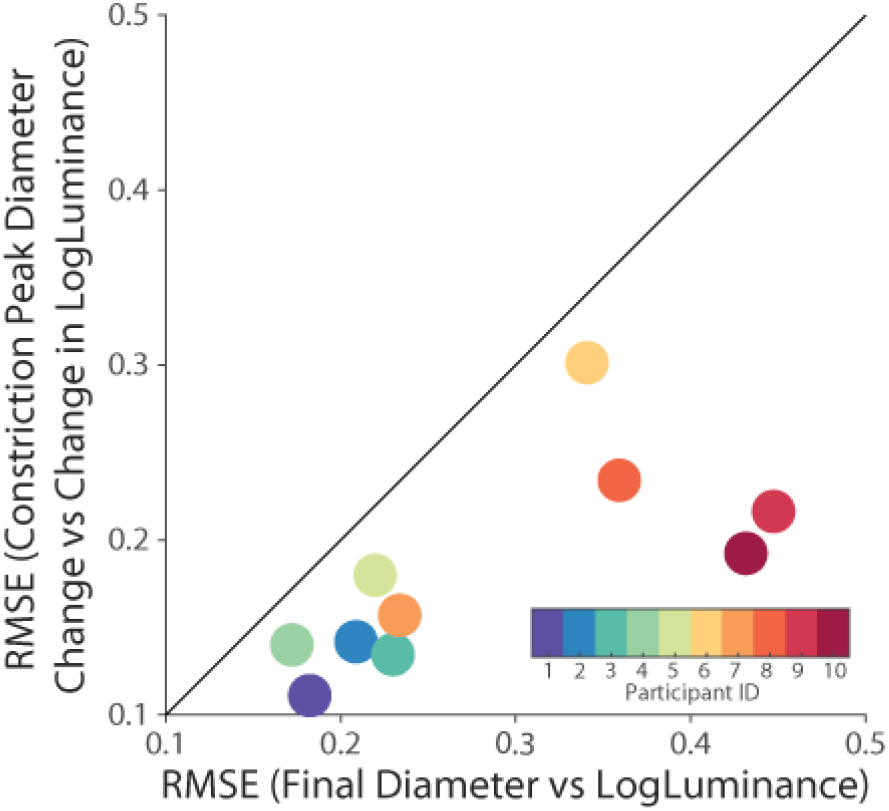
Within participant trial-by-trial variability in pupil metrics. The root-mean-squared errors of curve fits describing pupil metrics as a function of log-luminance are plotted for each participant. The unity line is illustrated in solid black. For each participant, variability in final pupil diameter (mm) as a function of log-luminance is consistently larger than variability in peak diameter change (mm) as a function of change in log-luminance in constriction trials (i.e. with dynamic overshoot).

We evaluated the covariability in transient dynamics for each participant. Figure 9 illustrates the relationship between peak velocity and peak diameter change. We observed qualitatively different trends for constriction vs dilation responses, but no longer observed the dramatic variability between participants apparent when considering the transient dynamics metrics as a function of the stimulus. We describe this highly stereotyped relationship across participants as the “Main Sequence”.

**Figure 9.**
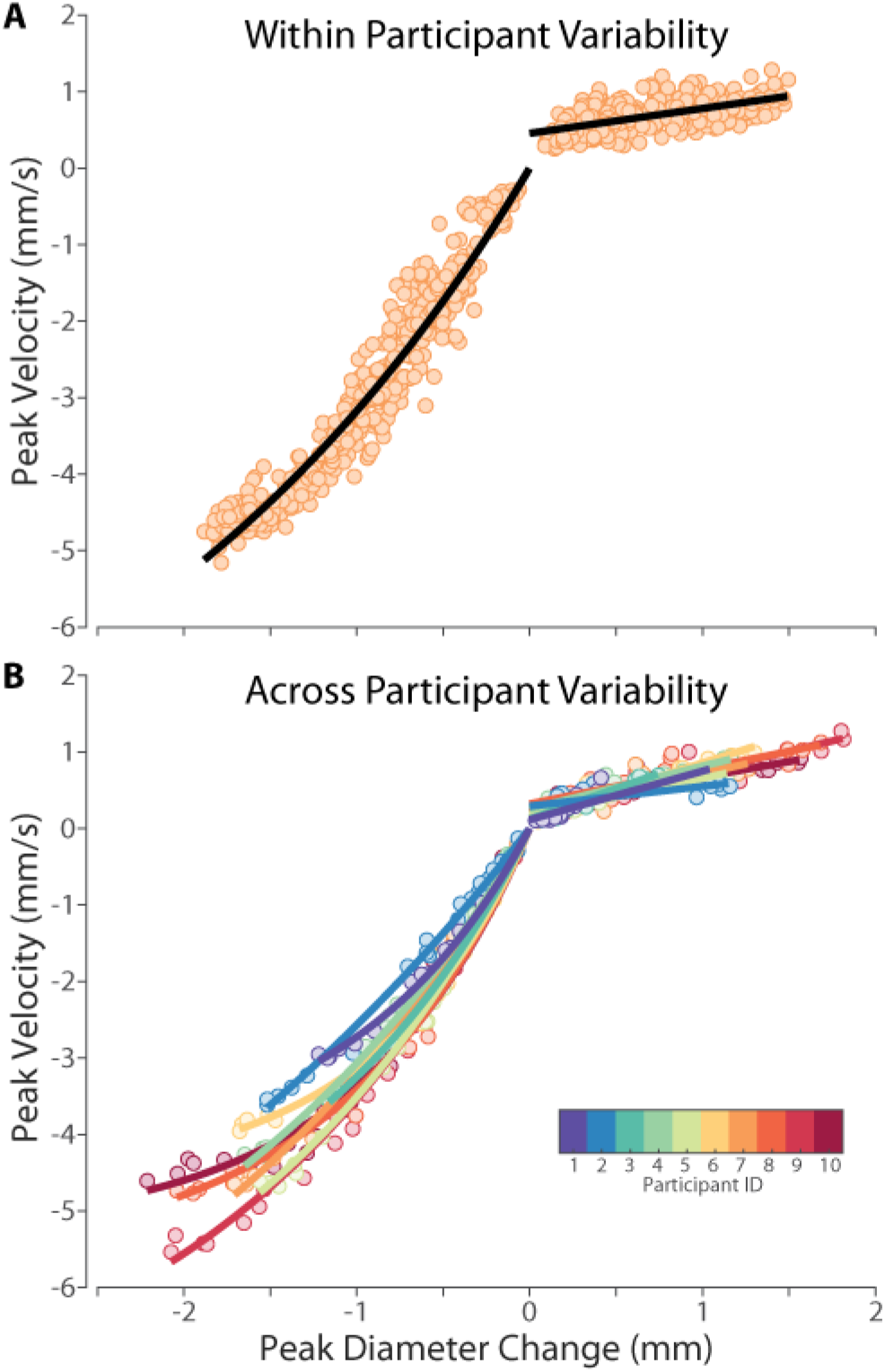
Human Pupil Main Sequence: Peak velocity vs peak change in pupil diameter. Dilation responses were well characterized by a linear function (y = a_1_*x + a_2_), except for small values where data tends towards 0 instead of a_2_. Exclusion of y-intercept parameter leads to unrealistic slope estimates. Constriction responses were better characterized by an exponential decay function, y = a_1_*(exp(x/a_2_) -1), capturing the saturation of peak velocity in large constriction responses. A) Trial-by-trial peak velocity vs peak diameter change for a single participant. B) Average peak velocity vs peak diameter change for each (initial, final) luminance condition for each participant.

## Discussion

The purpose of this study was to evaluate the dynamics of luminance evoked pupil responses in humans. We recorded participants pupil size during step changes in greyscale luminance during visual fixation. We observed that pupil diameter was linearly correlated with the logarithm of the luminance of the visual stimulus. Furthermore, we found correlated variability between pupil gain and pupil size across participants. We found high across-participant variability when considering transient dynamics metrics as a function of stimulus luminance, but a highly stereotyped “main sequence” relationship between peak velocity and peak diameter change across all participants. We first consider the limitations of the study, then discuss nonlinearities, variability, and the pupil main sequence. Finally, we consider the importance of these observations for the field moving forward.

### Limitations

First, our experimental set-up was designed to be representative of typical research lab environments, and therefore limited in the range of luminance values and the size of visual stimuli compared to natural conditions. As a result, pupil behaviour at the physiological extremes (both small and large pupil sizes) may not be captured in our data. While this limits the generalizability of our findings across environmental conditions, we believe this study is useful in characterizing pupil dynamics in the typical lab environments that pupillography will likely be employed in for research or clinical purposes.

Second, the behavioural task itself is quite unnatural (i.e., prolonged periods of visual fixation while viewing greyscale luminance changes while seated in a head and chin rest). In natural human environments, visual luminance typically varies across the visual field and over time and humans make eye movements (∼3 saccades/s), which will change the retinal input over time. Furthermore, we did not consider or control for the light-adaptation state of our participants, whose long term (> 1 hour) luminance history was unknown. Finally, our task used monochrome greyscale luminance, while the real world contains chromatic light that may differentially impact pupil responses (Barbur, 2004; Gamlin et al., 1998). Thus, our study does not assess within-participant variability due to light adaptation, the contribution of spatial and chromatic luminance variation to pupil responses, nor potential interactions between pupil responses and eye movements.

Third, the participants in our experiment were not representative of the full diversity of humans, nor did our experiment explore the full diversity of within-participant variability. Nevertheless, this demographic is a valuable starting point for future research systematically testing the influence of age and clinical pathology on pupil dynamics. This group may be a typical ‘control’ group, yet variability of pupil responses in young, healthy is underappreciated in literature. Thus, further research is required to systematically evaluate the regularity of the main sequence relationship and deviations from the main sequence.

### General Discussion

We have quantified several nonlinearities in pupil response metrics. First, steady-state pupil diameter is a nonlinear function of luminance. This relationship can be linearized by considering the logarithm of luminance, i.e. changes in steady-state pupil diameter depend on the ratio of luminance change (Clarke et al., 2003; Ellis, 1981; Gamlin et al., 1995; Pong and Fuchs, 2000a, 2000b; Watson and Yellott, 2012). This relationship should be accounted for when performing analysis that assume/require linearity in stimulus-response relationships (e.g., by first log-transforming stimulus luminance). Second, peak diameter change and peak velocity are still nonlinear functions of log-luminance for constriction (Bremner, 2012; Ellis, 1981) but not dilation. Similarly, the constriction side of the pupil main sequence shows a nonlinear relationship. Thus, the nonlinear relationships in these metrics should be considered to avoid inappropriate curve-fitting or statistical analysis methods that would lead to biased inferences.

Our data demonstrates a high degree of across-participant variability among young, healthy adults. The across-participant variability can be seen in all stimulus-response relationships. However, we found correlated variability between overall pupil size and pupil gain across participants. Thus, participants with overall larger pupils also showed larger diameter changes per unit log-luminance change. This type of across-participant variability cautions the use of baseline normalization (see also Mathôt et al., 2018; Mathôt and Vilotijević, 2023 for other cautions about normalizing pupil data). For example, if average pupil diameter varies between groups, the relationship between pupil size and pupil gain provides the *a priori* expectation that the group with the larger pupils will also show larger responses. A potential strategy could be to control for average pupil size during participant selection (e.g. finding pupil size matched controls). Another potential strategy could be to normalize each participant’s pupil responses by their specific pupil gain before making inferences about stimulus strength or effect size (for evaluating differences in pupil responses due to some intervention).

We also observed within-participant variability, described in terms of trial-by-trial variability in pupil response metrics. Specifically, we observed greater variability in final pupil size compared to transient constriction peak amplitude. The timing of the transient constriction responses is relatively early and time-locked to the luminance, which suggests an accumulation of variability from the time of constriction peak amplitude until the time of final diameter calculation. The sources of trial-by-trial variability remain beyond the scope of this study and merit further investigation.

Despite the variability described above, we found a well conserved pupil main sequence: a non-linear lawful relationship between peak velocity and peak diameter change for trial-by-trial luminance evoked pupil responses across all participants. The pupil main sequence does not consider the sensory inputs (i.e. the ongoing luminance values/changes) that produce trial-by-trial pupil responses, instead describing covariability in motor parameters of the transient pupil response. This analysis of motor covariability complements typical stimulus-response style pupil analyses and our previous discussion of across participant variability in terms of stimulus-response parameters, e.g. across participant variability in pupil gain. Similar to the saccade main sequence (Bahill et al., 1975), the structured covariability observed in the pupil main sequence could be informative for modelling the pupil control system. This relationship seems compatible with a pulse-step controller (Robinson, 1973; Semmlow and Stark, 1971; Usui and Hirata, 1995), particularly for constriction responses where large saturating peak velocities are observed. This hypothesis is compatible with neuronal responses in the primate pupillomotor control pathway, including the burst-tonic activity of intrinsically photosensitive retinal ganglion cells (Dacey et al., 2005) and pretectal olivary neurons (Pong and Fuchs, 2000b). On the other hand, the asymmetries between constriction and dilation dynamics suggest potential differences in their underlying control (Longtin and Milton, 1989; Semmlow and Stark, 1971). Continued research is required to synthesize a model of pupil control that can account for the main sequence.

The main sequence relationship can also be used in future research investigating abnormalities or variability in pupil behaviour by further evaluating the regularity of the main sequence and the conditions resulting in deviations from this stereotypical motor covariability. For example, pupil responses during accommodation seem to show similar covariability between peak velocity and peak amplitude change, including the differences between constriction and dilation (Kasthurirangan and Glasser, 2005), although the relationships and interactions between pupil responses to light, accommodation, and vergence merits further investigation. On the other hand, it is not clear if the main sequence motor covariability persists for cognitive-related pupil behaviours. For example, cognitive modulations of the pupillary light response (Binda and Murray, 2015; Ebitz and Platt, 2015; Mathôt et al., 2013; Steinhauer et al., 2000; Wang et al., 2018a) may induce variability along the main sequence through changes in pupil gain or induce variability orthogonal to the main sequence through changes in motor programming – similar to the relationship between subjective value and variability in saccade peak velocity orthogonal to the saccade main sequence (Reppert et al., 2015). Thus, the main sequence could be used as a research tool to investigate pupil behaviour and the neural control of pupil dynamics.

## Conclusions

We evaluated luminance evoked pupil dynamics in young healthy adult humans. We describe nonlinearities in pupil response metrics, variability within and across participants, and the stereotyped pupil main sequence relationship describing motor covariability in pupil dynamics. Our study highlights the need for more research to evaluate within and across participant variability and their sources. The main sequence should provide new avenues for investigating and modelling pupil variability and the neural control of pupil dynamics, including investigating the effects of light adaptation, aging, the occurrence and progression of neurological disorders, and pharmacological interventions.

## Appendix

We compared the diameter of left and right pupils for each subject and session, which is presented in Table 1. When considering a linear model with a bias term, most subjects had near unity slope and a bias within the range of physiological anisocoria (<0.4 mm, not considered a clinical concern) (Steck et al., 2018; Wang et al., 2018b).

**Table 1.** Bilateral pupil response linear regression results.

| Subj# | Session# | Right Pupil = A*Left pupil + B |  |  |  |  |  |
| --- | --- | --- | --- | --- | --- | --- | --- |
|  |  | A | Confidence Interval |  | B (mm) | Confidence Interval (mm) |  |
| 1 | 1 | 0.993 | 0.986 | 1.000 | 0.117 | 0.092 | 0.142 |
| 1 | 2 | 0.953 | 0.941 | 0.965 | 0.090 | 0.041 | 0.138 |
| 2 | 1 | 1.001 | 0.986 | 1.017 | 0.269 | 0.202 | 0.336 |
| 2 | 2 | 1.039 | 1.023 | 1.055 | 0.216 | 0.155 | 0.277 |
| 3 | 1 | 0.919 | 0.903 | 0.935 | 0.340 | 0.291 | 0.390 |
| 3 | 2 | 0.950 | 0.937 | 0.963 | 0.090 | 0.048 | 0.132 |
| 4 | 1 | 0.715 | 0.691 | 0.740 | 0.471 | 0.397 | 0.545 |
| 4 | 2 | 0.795 | 0.777 | 0.814 | 0.420 | 0.369 | 0.470 |
| 5 | 1 | 0.903 | 0.893 | 0.913 | 0.146 | 0.119 | 0.174 |
| 5 | 2 | 0.960 | 0.948 | 0.971 | 0.105 | 0.075 | 0.136 |
| 6 | 1 | 0.961 | 0.946 | 0.976 | 0.049 | 0.006 | 0.091 |
| 6 | 2 | 0.898 | 0.886 | 0.910 | 0.170 | 0.134 | 0.206 |
| 7 | 1 | 1.087 | 1.065 | 1.108 | -0.158 | -0.210 | -0.106 |
| 7 | 2 | 1.026 | 1.005 | 1.046 | -0.054 | -0.105 | -0.003 |
| 8 | 1 | 0.907 | 0.896 | 0.917 | 0.239 | 0.205 | 0.273 |
| 8 | 2 | 0.915 | 0.902 | 0.927 | 0.180 | 0.143 | 0.217 |
| 9 | 1 | 1.003 | 0.992 | 1.014 | 0.013 | -0.011 | 0.037 |
| 9 | 2 | 1.049 | 1.037 | 1.060 | -0.039 | -0.064 | -0.014 |
| 10 | 1 | 1.053 | 1.038 | 1.068 | -0.060 | -0.095 | -0.026 |
| 10 | 2 | 1.041 | 1.014 | 1.068 | 0.078 | 0.023 | 0.134 |

## Notes

https://osf.io/vrch6/

https://github.com/BlohmLab/PupilMainSequence

### Competing Interest Statement

The authors have declared no competing interest.

